# Back-to-the-future motion analysis using machine intelligence predicts the potential risk of mood disorders

**DOI:** 10.1101/2024.11.12.623133

**Authors:** Wongyo Jung, Junesu Lee, Sujin Chae, Daegun Kim, Daesoo Kim

## Abstract

Motion encodes emotional value. Decoding the hidden information within the complexity and randomness of motion may aid in the early diagnosis and prevention of mood disorders. In this study, we utilized machine intelligence to predict the risk of mood disorders by analyzing random motion patterns in mice. The support vector machine, trained on motion data collected before chronic social defeat stress (CSDS), predicted a significant correlation between these data and the degree of CSDS-induced social avoidance. Unsupervised clustering revealed key motion biomarkers of stress susceptibility, accounting for approximately 70% of the onset of social avoidance in a new group of mice before exposure to CSDS. These findings suggest that motion analysis using machine intelligence could provide a non-invasive approach for predicting the risk of mood disorders.

## INTRODUCTION

Mood disorders develop unpredictably owing to complex interactions between genetic and environmental factors. Why an individual only develops mood disorders after experiencing chronic stress is a long-standing critical question in the field. Studies using inbred mice with the same genetic background have demonstrated differential epigenetic changes in the brain in response to chronic stress between susceptible and resilient mice (Tsankova et al., 2006; Renthal et al., 2007; Hunter et al., 2009; Wilkinson et al., 2009; Covington et al., 2011; Golden et al., 2013; Dudek KA et al., 2020). However, when susceptibility to stress occurs – before, during, or after exposure to stress – is not yet clear.

One of the primary challenges in resolving this question is that susceptible and resilient animal subjects often exhibit strikingly similar behaviors during freely moving states, particularly in open-field tests (OFTs) before and after stress experiences (Krishnan et al., 2007; Liu et al., 2017; Murra et al., 2022). This phenomenon underscores the difficulty of studying stress susceptibility in the absence of stress exposure and underscores the need to develop sensitive methods for identifying biomarkers associated with stress susceptibility that allow distinguishing been susceptible and resilient subjects. Importantly, if such methods are expected to analyze further causal relationships related to the future onset of mood disorders, they must avoid sacrificing animals or causing further stress.

Recent advances in machine intelligence have revolutionized the field of computational ethology, enabling automated measurement and analysis of behaviors, in the process uncovering hidden information in behavioral patterns that have remained elusive using traditional methods (Mathis et al., 2018; Pereira et al., 2019; Dolensek, 2020; Marshall et al., 2021). In this context, our study presents a novel approach for tackling the dilemma of studying stress susceptibility. Leveraging the AVATAR (AI-Vision for Action Translation And Reconstruction) system, which enables markerless motion sequencing by detecting the 3D coordinates of key body points (Kim et al., 2021), we introduce a pipeline for identifying motion biomarkers by comparing past motion data between groups of freely moving animals classified based on results of subsequent chronic social defeat stress (CSDS).

## RESULTS

### “Back-to-the-future” motion sequencing for the analysis of stress susceptibility prior to experiencing chronic social stress

To compare motion patterns between mice that are susceptible and resilient to chronic stress before they experience stress events, we designed an experimental procedure that we call “Back-to-the-Future Motion Analysis” (Fig. 1). In this procedure, we initially assessed the behavior of freely moving C57BL/6J mice by conducting an open-field test (OFT). We then exposed the mice to social stressors for 10 minutes daily over 10 days to gauge stress susceptibility. At the end of the chronic social defeat stress (CSDS) protocol, we categorized the mice based on their social preference index (SI) during a novel encounter as follows: susceptible, with a higher social avoidance index (SI < 1, n = 28), and resilient, lacking social avoidance (SI ≥ 1, n = 15) (Extended Data Fig. 1a), as described previously (Krishnan, Vaishnav et al., 2007; Chae S et al., 2021). Finally, we compared the motion data obtained before CSDS between the two groups of freely moving mice. We found no differences in conventional parameters, such as total distance moved and exploration of the center zone, consistent with previous findings (Krishnan et al., 2007; Murra et al., 2022) (Extended Data Fig. 1b–e).

**Fig. 1:**
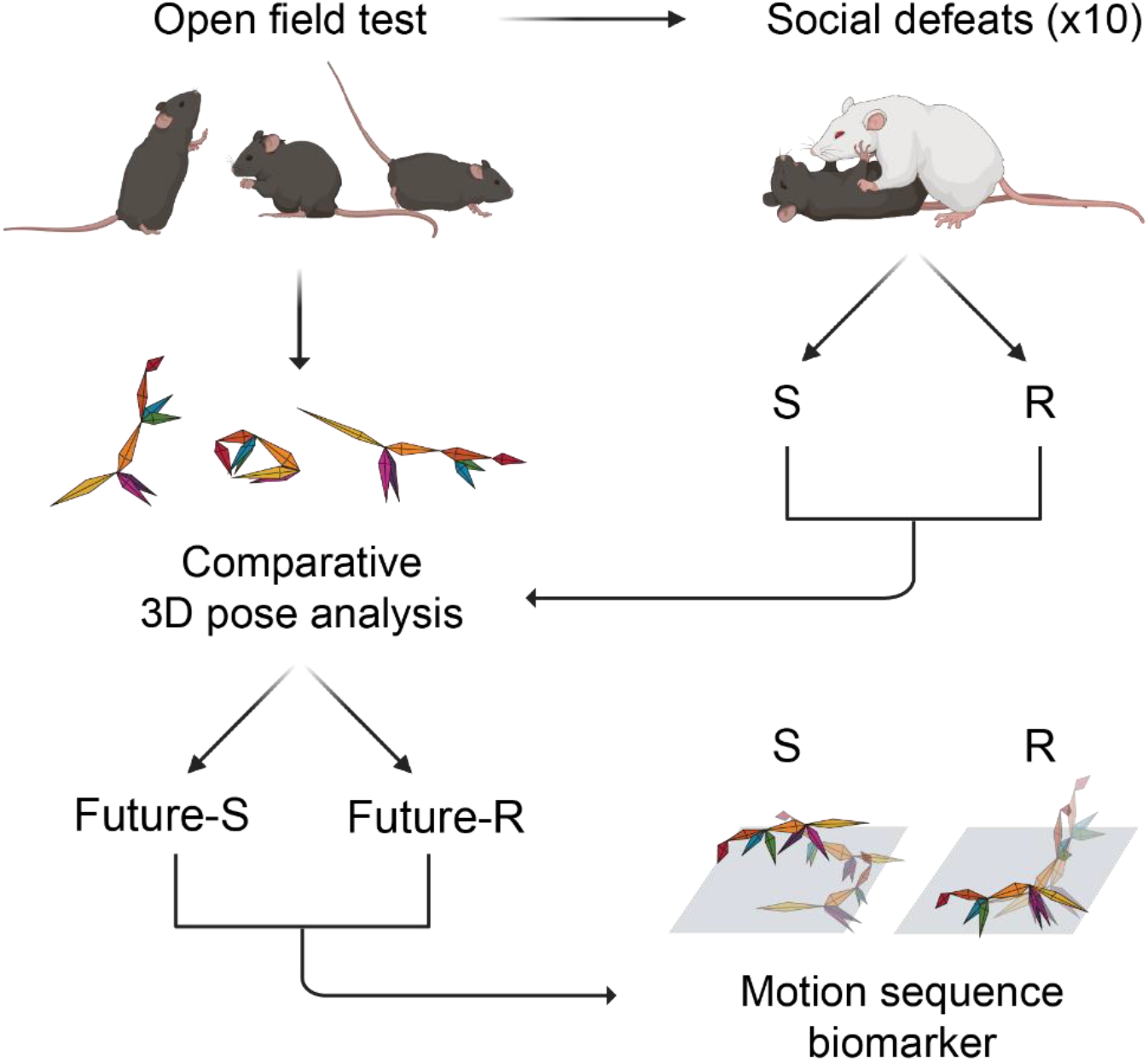
Workflow of stress susceptibility prediction before chronic stress. Schematic illustration showing analysis pipeline. Behaviors in the video are transformed into 3D coordinates of each body point of a mouse; mice are then classified into susceptible (S) versus resilient (R) groups according to a social interaction score measured after chronic social defeat stress. Behavioral patterns in future susceptible and resilient individuals are distinguished through comparative analysis using dimensional reduction and clustering techniques. Finally, sequential motion biomarkers for depression diagnosis are extracted.

### Susceptible mice show different freely moving behavior from resilient mice

To comprehensively analyze freely moving behaviors in the OFT measured before CSDS, we utilized the deep–learning–based 3D pose estimation system, AVATAR (Kim et al., 2021), which transforms video motion data into a numerical motion sequence comprising 3D coordinates (*x, y, z*) of nine key body points (nose, head, body center, anus, right and left front limbs, right and left hind limbs, and tail tip) at a 10-millisecond resolution. We then employed these motion data to compare the two groups of mice and identify motion biomarkers associated with stress susceptibility.

We trained a support vector machine (SVM) using motion sequence data of varying lengths (1–10 seconds × 3 coordinates × 9 body points) and evaluated its decoding performance on a test dataset (Fig. 2a). The SVM decoder exhibited a decoding accuracy ranging from 50% to 70%, showing no proportional correlation with data length (Fig. 2b). Consequently, we chose 4-second motion sequences for training the SVM. Employing this length unit, the SVM successfully discriminated between the two groups of mice (Fig. 2c). We further confirmed that other supervised classifiers, including naive Bayesian and random forest classifiers, showed similar decoding accuracy (Fig. 2c). Subsequently, we classified the future susceptibility of mice by averaging scores from all motion sequences for each mouse. Classification using the average score from the 4-second motion sequence data demonstrated higher decoding accuracy in the test dataset than expected in the random-shuffled dataset, demonstrating a robust correlation with individual SI values in new datasets (61.17% vs. 49.56%; Fig. 2d, e).

**Fig. 2:**
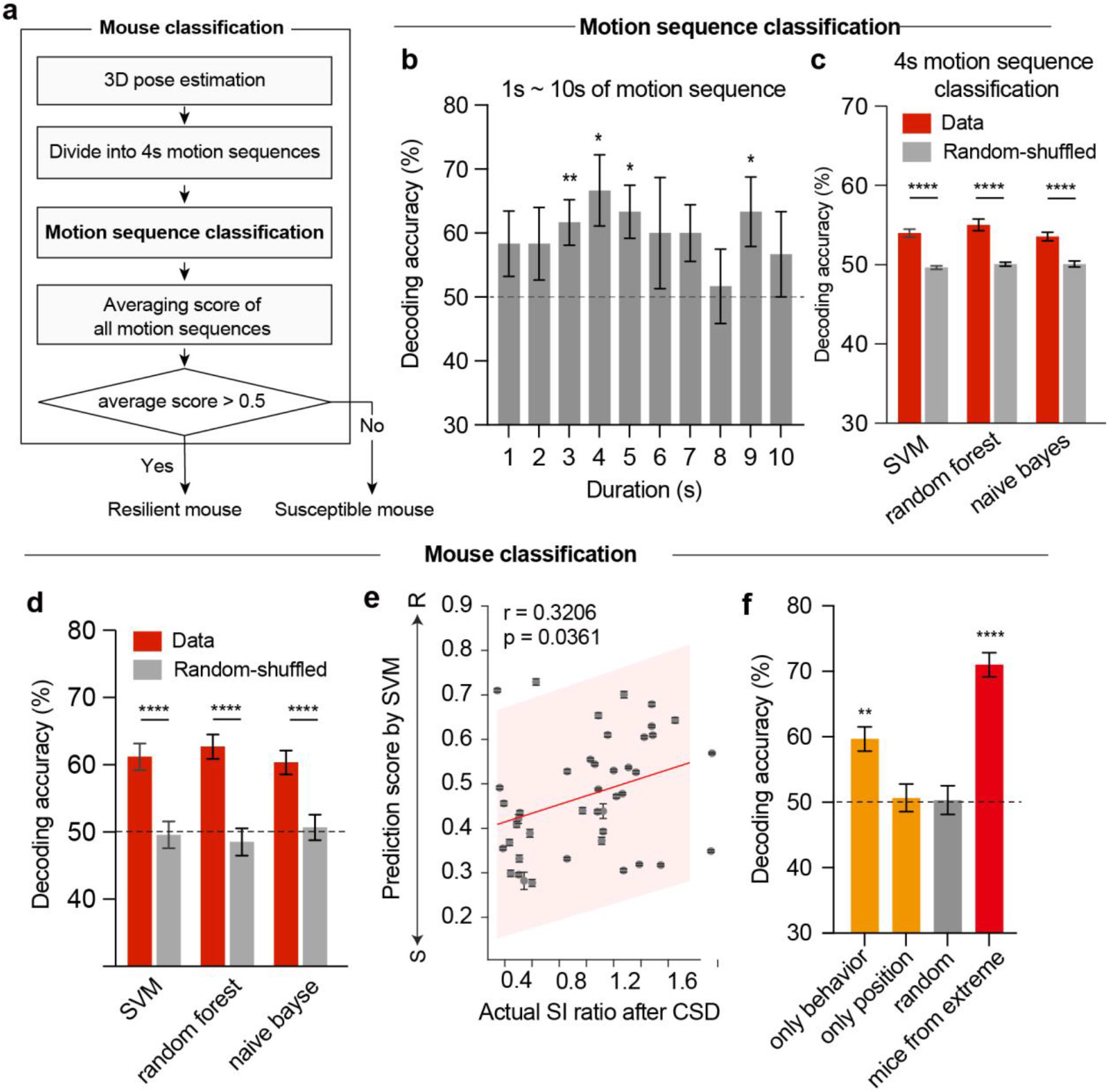
Classification of future resilience and susceptibility in mice based on open-field behavior. a, Analysis pipeline. RGB video from five multi-view cameras is converted into 3D pose coordinates using AVATARnet, and the continuous coordinates are divided into motion sequences. Machine-learning classifiers are trained on the motion sequences to classify resilient and susceptible motion sequences (motion sequence classification). All scores of motion sequences for each mouse are then averaged, and mice are segregated into resilient and susceptible categories using a score threshold of 0.5 (mouse classification). b, Decoding accuracy of the SVM decoder across varying durations of motion sequence (1 s, 58.33%; 2 s, 58.33%; 3 s, 61.67%; 4 s, 66.67%; 5 s, 63.33%; 6 s, 60%; 7 s, 60%; 8 s, 51,67%; 9 s, 63.33%; 10 s, 56,67%) during 10 random iterations. A total of 15 resilient and 15 susceptible mice were randomly selected for each iteration. The dashed black line marks the 50% (chance) level. A one-sample t-test was used to compare each decoding accuracy with 50%. c, Comparison of 4-second motion sequence decoded from three machine-learning classifiers using the hold-out cross-validation method with 100 repetitions. Each decoding accuracy was compared with the random-shuffled dataset using a t-test. (SVM: 53.98%, random forest: 55.01%, naive Bayse: 53.54% for dataset; SVM: 49.64%, random forest: 50.07%, naive Bayse: 50.09% for shuffled data). d, Comparison of mouse classifications from three machine-learning classifiers using hold-out cross-validation with 100 repetitions. Each decoding accuracy was compared with the random-shuffled dataset using a t-test. (SVM: 61.17%, random forest: 62.67%, naive Bayse: 60.33% for dataset; SVM: 49.56%, random forest: 48.5%, naive Bayse: 50.67% for shuffled data). e, Scatter plot of predicted and actual social interaction scores in all mice from the test dataset. The predicted score (1 for resilient, 0 for susceptible) for each mouse is an average of all predicted motion sequence scores during the 400-second OFT. r, correlation coefficient. f, Comparison of decoding accuracy of the SVM, trained with different data features. The test dataset was randomly selected using hold-out validation for 100 iterations (behavior only, 59.67%; position only, 50.67%; random, 50.33%; mice at the extreme, 71%). Data are presented as means± s.e.m. (**P < 0.01, ***P < 0.001, ****P < 0.0001; one-way ANOVA with Dunnett’s post hoc comparison).

We further processed the motion data, dividing them into motion sequence data (relative 3D coordinates of 9 body parts from the fixed position) and positional data (3D coordinates of the body’s center point). We then trained the SVM decoder on these datasets and assessed each decoding performance. The SVM had no predictive capacity when trained solely on positional data (50.67%; Fig. 2f) – a key parameter in conventional analysis – as previously described (Krishnan et al., 2007; Pena et al., 2017; Willmore et al., 2022) and shown in our study (Extended Data Fig. 1d). In contrast, when trained on motion sequence data without positional information, the SVM decoder showed higher accuracy than that trained with randomly labeled data (59.67% vs. 50.33%; Fig. 2f). In addition, when trained on motion sequence data obtained from extremely susceptible mice (lowest SI rankers, n = 8) and resilient rankers (highest SI rankers, n = 8), the prediction accuracy increased to 71%. These findings support the existence of specific motion sequences associated with stress susceptibility even before CSDS.

### Extraction of motion sequences associated with stress susceptibility

Next, we sought to identify specific motion sequences linked to stress susceptibility, focusing on those with a predicted higher incidence in susceptible mice compared with resilient mice, with the goal of predicting the onset of CSDS-induced social fear in new test groups (Fig. 3).To capture motion sequences that differentiate highly susceptible and resilient groups, we initially aligned motion sequences so that they commenced in the same quadrant of the chamber, thereby minimizing positional information bias (Fig. 3a). We then selected motion sequences correctly classified by the decoder and conducted supervised dimensionality reduction using partial least squares (PLS), aiming to find the lowest dimension that effectively separated the motion sequences of resilient and susceptible mice (Fig. 3b).

**Fig. 3:**
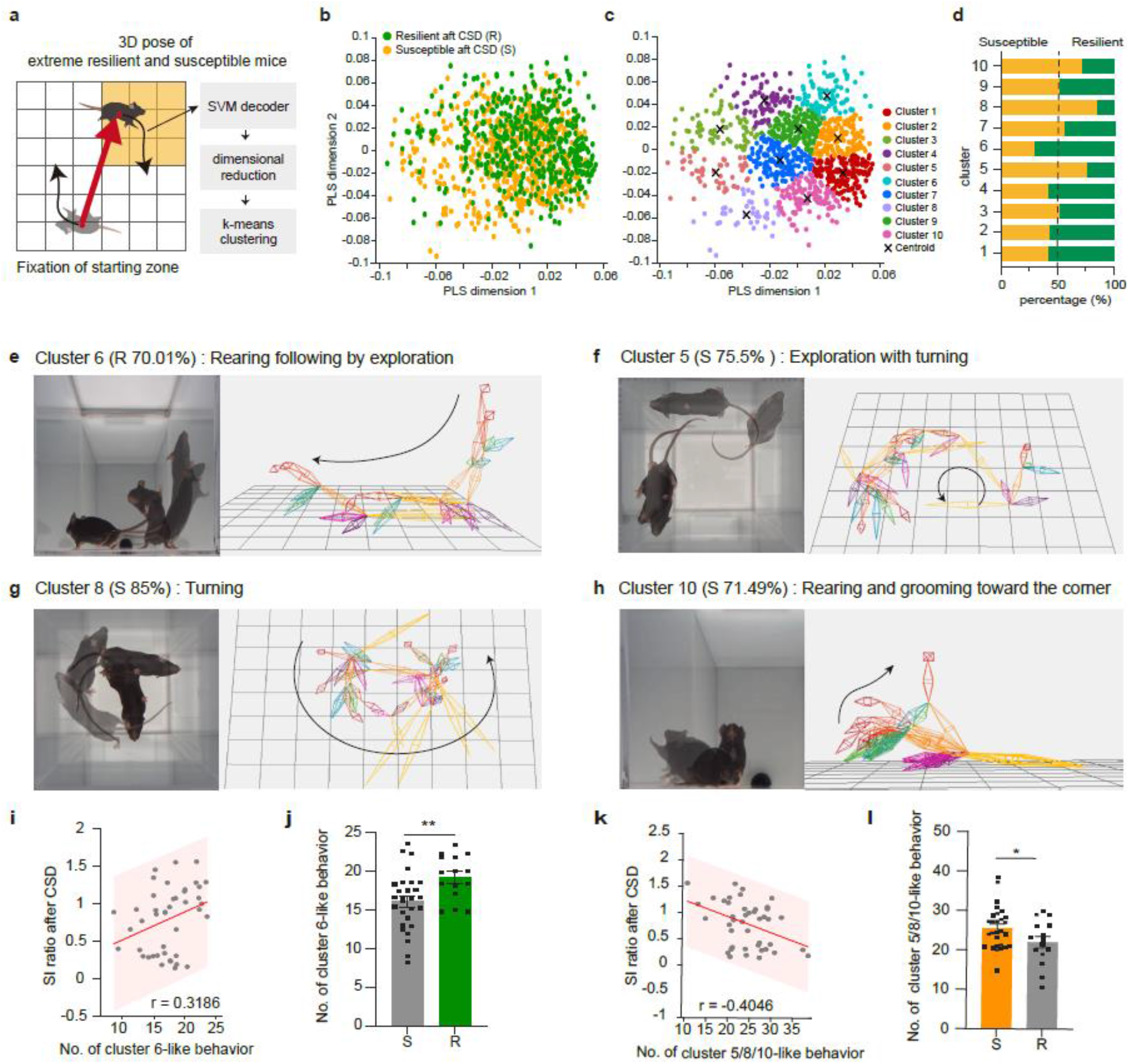
Dimensional reduction reveals characteristic behavior features in resilient and susceptible mice before chronic social stress. a, Analysis schematics for feature extraction using dimensional reduction and unsupervised k-means clustering. Data were pre-processed to have the same starting zone and all motion sequences starting from the first quadrant of the *x-y* axis. b, PLS visualization of motion sequences correctly classified by the SVM decoder (n = 1041 data points, variance explained by first dimension: 2.86%, by second dimension: 1.55%). c, K-means clustering of data points in the PLS visualization. d, Susceptible/resilient data points ratio in each cluster in (c). e–h, Representative motion sequences of cluster 6 (e), cluster 5 (f), cluster 8 (g), and cluster 10 (h). i,k, Scatter plot of social interaction scores and number of cluster 6-like (i) and cluster 5/8/10-like (k) behaviors in each mouse. j,l, Number of cluster 6-like (j; P = 0.0083) and cluster 5/8/10-like (l; P = 0.0499) behaviors in susceptible and resilient mice. Data are presented as means± s.e.m. (*P < 0.5, **P < 0.01; unpaired t-test). r, Pearson’s correlation coefficient.

Notably, the motion sequences of the two groups exhibited distinct distributions in the PLS dimensional space, with sequences from susceptible mice displaying a broader distribution that did not overlap with that of resilient mice. After embedding the motion sequences into this reduced dimension, we applied unsupervised k-means clustering to group behaviors based on their proximity (Fig. 3c). This analysis identified 10 clusters, each comprising various combinations of motion sequences from resilient and susceptible mice. Notably, four of these clusters (clusters 5, 6, 8, and 10) showed a dominant proportion (>70%) of one group relative to the other (Fig. 3d).

To gain insights into each behavior cluster, we reconstructed representative motion sequences from each cluster, converting them into 3D coordinates (Extended Data Fig. 2, Supplementary Video, and Methods). Resilient mice showed more frequent rearing behavior followed by exploration (cluster 6), a pattern different from those of other rearing clusters, such as cluster-1, with rearing then crouching; cluster-2, with more prolonged rearing; and cluster-4, with rearing and moving to the corner (Fig. 3e and Extended Data Fig. 2). In contrast, susceptible mice showed three dominant clusters (5, 8, and 10); clusters 5 and 8 featured similar behaviors (turning followed by exploration), whereas cluster 10 exhibited a higher frequency of grooming behaviors in the corner (Fig. 3f, g, and h). These behaviors differed from those observed in other clusters (Extended Data Fig. 2).

Next, we trained the SVM decoder on specific motion sequences one by one and tested whether it distinguished the two groups of mice with extreme SI values in a freely moving state in the OFT. Interestingly, cluster-6 exhibited a significant positive correlation with the SI ratio after CSDS (Fig. 3i). Cluster 6-like motion sequences occurred with significantly different frequencies between susceptible and resilient mice, with the susceptible group displaying these sequences more frequently (P = 0.0083; Fig. 3j). Although clusters 5, 8, and 10 were not individually correlated with SI value (Extended Data Fig. 3), an analysis using all three of these clusters showed a strong correlation (Fig. 3k) and different incidences between resilient and susceptible mice (P = 0.0499; Fig. 3l).

### Validation of motion sequence biomarkers for prediction of stress susceptibility

Our investigation primarily centered on determining whether dominant behavioral features from one particular group – susceptible or resilient – could serve as dependable predictors of the SI ratio following CSDS. This inquiry was driven by a discernable correlation between the prevalence of dominant behavioral features in one group and the SI ratio, a relationship initially established based on data from the extreme groups (Fig. 3).

To explore the universal applicability of these behavioral predictors across all datasets, including a broader population of mice not originally characterized for these behaviors, we employed a logistic generalized linear model (GLM) (Fig. 4a). This model utilized the count of susceptible-dominant and resilient-dominant cluster-like behaviors observed during the OFT as predictor variables. The goal in applying this model was to forge a mathematical relationship between the exhibited behaviors and subsequent SI ratios after CSDS, using both its classification accuracy and correlation with actual SI ratios as a gauge of its effectiveness. The GLM showed promising enhancement in classification accuracy, successfully differentiating between resilient and susceptible mice with 67.5% accuracy when trained on the selected behavioral features (Fig. 4b). This marked an improvement from a baseline accuracy of 61.17% (Fig. 2d). Notably, the model’s predicted scores also closely mirrored the distribution of the actual SI ratios (Fig. 4c) and showcased a stronger positive correlation with the SI ratio than was achieved using the SVM decoder (0.4059 vs. 0.3206, respectively) (Fig. 4d and Fig. 2d). Moreover, the GLM predicted resilient mice with slightly higher accuracy compared with susceptible mice, underscoring a potentially more significant role of resilient-like behaviors in predicting post-stress sociability levels (Fig. 4e).

**Fig. 4:**
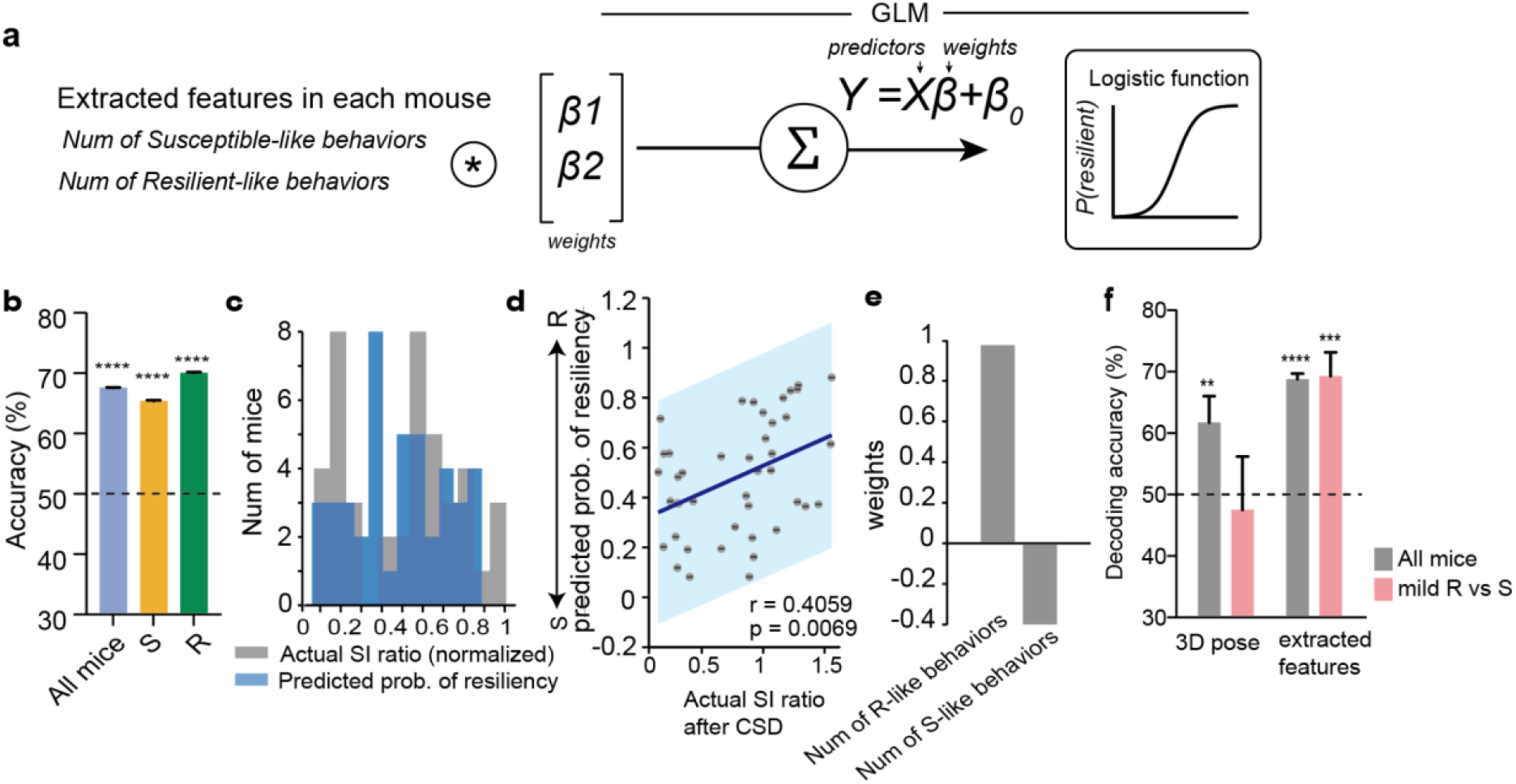
Decoded characteristic behavior is correlated with the susceptibility of mice. a, Schematic depiction of the generalized linear model (GLM) for encoding the relationship between the characteristic features and social interaction score. The binary distribution and logistic link function of the GLM were used for binary classification (resilient vs. susceptible). b, Predicted accuracy of GLM classification of resilient and susceptible mice (Average: all mice, 67.5%; S, 65.39%, R, 70.07%; ****P < 0.0001; unpaired t-test,) c, Scatter plot of predicted resilient probability and social interaction score. R, Pearson’s correlation coefficient. d, Fitted GLM weights for each parameter. The magnitude of each weight represents the parameter’s relative importance to the regression of the SI score, whereas the sign of each weight indicates the direction of each parameter’s correlation with the SI score. e, Comparison of decoding accuracy between the SVM decoder trained on 3D pose data and selected behavioral features. The pink/blue bar accurately calculates extreme/mild resilient and susceptible mice. The dashed line represents the 50% (chance) level. f, Quantification of the accuracy of the SVM trained on all mice and mild resilient and susceptible mice in 10 times repeated random sample selection (**P < 0.01, ****P < 0.0001 vs. chance (50%); one sample t-test).

To more rigorously validate the applicability of the identified features and understand the characteristics of the mild group, comprising mice with middle-range SI ratios (0.2875 to 1.1882), we utilized an SVM decoder for classification based on SI ratio displayed by the mice. While the SVM decoder yielded an accuracy of 47.5% when trained solely on 3D pose data from the mild group, it demonstrated enhanced performance, reaching an accuracy of 66.88% when trained on the extracted behavioral features (Fig. 4f).

This enhanced accuracy in classification is a testament to the steadfastness and reliability of the identified behavioral features. Their ability to accommodate variations in individual mouse behaviors, together with their adaptability to diverse decoding methods, highlights the robustness of these behavioral features. Collectively, these attributes collectively emphasize the broad applicability and potential utility of these behavioral signatures, underlining their relevance and potential impact on subsequent research initiatives.

## DISCUSSION

Understanding the intricate and seemingly random movements of animals during their free-moving states presents a challenging to researchers, as it involves identifying motion biomarkers that encode specific information beyond the capacity of human cognition. In this study, we leveraged computational ethology technology, as previously introduced (Anderson et al., 2014; Mathis et al., 2018; Kim et al., 2022), to uncover motion biomarkers that can predict susceptibility to future mood disorders resulting from chronic stress exposure (Fig. 1).

The complex behavioral “dance” of animals in a freely moving state holds valuable insights into their potential vulnerability to chronic stress, as depicted in Figure 2. These insights are encapsulated within distinct action patterns characterized by the spatial coordination of various body parts, as illustrated in Figure 3. Importantly, they effectively serve as a prelude to an individual’s risk of succumbing to chronic stress, as shown in Figure 4d.

Prior research has extensively explored the consequences of chronic social defeat stress in terms of neurological, endocrine, and behavioral outcomes (Krishnan et al., 2007; Milic et al., 2021), but the enigmatic question of whether intrinsic factors contribute to these variations in mice before their exposure to stress has remained unanswered. Analyzing behavioral traits in the absence of social stress presents unique challenges. Unlike stressful events with distinct time points for analysis, spontaneous behaviors that occur in the absence of explicit stress stimuli lack such temporal markers.

To bridge this gap, we introduced data-driven and machine-learning approaches, enabling the identification of behavioral features associated with susceptibility and resilience in mice even before they encounter social stress. Remarkably, the absence of discernible differences in anxiety-like behavior between resilient and susceptible mice before chronic stress exposure (Extended Data Fig. 1) suggests that the observed behavioral disparities are not attributable to mild stress incurred during the OFT (Seibenhener et al., 2015). Instead, it is plausible that resilient and susceptible mice employ distinct exploration strategies or exhibit nuanced behavior patterns. In future studies, the application of machine intelligence approaches that combine other physiological factors, including biochemical and neurophysiological phenomena, with motion biomarkers may provide more precise prediction at the individual level.

Understanding the underlying neurological and physiological factors that govern these behavioral features is a compelling avenue for future exploration. Building upon prior findings that mice adhere to general behavioral principles characterized by specific stereotypical and patterned motion sequences (Tinbergen, 1951; Wiltschko et al., 2015; Markowitz et al., 2018; Marshall et al., 2021; Markowitz et al., 2023), we have demonstrated the utility of machine intelligence in finding additional traits hidden within random movements. This knowledge not only forms a framework for improving the quality and ethical aspects of animal experiments, but also paves the way for early diagnosis and prediction of neurological disorders through harnessing the immense potential of machine intelligence.

## Materials and Methods

### Mice

In each experiment, 53 adult male C57BL/6J mice (9–15 weeks old; Jackson Laboratories stock #000664) were utilized. Before being subjected to the chronic social defeat stress test, mice were communally housed in groups of four to six per cage. CD-1 retired breeder mice (>15 weeks old) served as the aggressor animals in the chronic social defeat stress paradigm. Mice were maintained under a 12-hour light-dark cycle and were provided food and water ad libitum. All procedures were conducted according to Korea Advanced Institute of Science and Technology (KAIST) Guidelines for the Care and Use of Laboratory Animals and were approved by the Institutional Animal Care and Use Committee (Protocol No. KA2020-65).

### Open field test

We employed the AVATAR studio for the OFT and recording, as described by Kim et al. (2022). A transparent chamber constructed from 5-mm–thick acrylic panels (200 mm x 200 mm x 300 mm) was placed at the center of the AVATAR studio, where mice were allowed to move freely. The spontaneous behavior of mice was captured by five high-speed cameras (BFS-U3-23S3C-C; FLIR Systems, Inc., Richmond, BC, Canada) installed at distinct viewpoints – all four sides and bottom. The cameras’ specific positions were adjusted based on their focal length.

Each wild-type mouse was recorded individually in the studio. After a 30-minute habituation period in the experimentation room, each mouse’s behavior was recorded by the five cameras for at least 400 seconds at a rate of 30 frames per second. Images from each of the five cameras were merged into a single composite image.

### Chronic social defeat model

Retired CD-1 breeder mice (>15 weeks old) were evaluated for aggression, as previously documented (Golden et al., 2011). Each C57BL/6J mouse, aged 11 weeks, was exposed to physical defeat involving an unfamiliar, aggressive CD-1 mouse. After an initial aggressive interaction, CD-1 and C57BL/6J mice were separated for 24 hours by a transparent acrylic partition with multiple holes, permitting sensory contact. Non-defeated control mice (i.e., those not exposed to a CD-1 mouse) were housed similarly and subjected to an equivalent rotation schedule. Following a 10-day social defeat period, all mice were housed individually.

### Social interaction test

The social avoidance test was conducted in a dimly lit room using an amber open-field box (42 × 42 cm) containing a removable iron mesh cage (10 × 6.5 × 42 cm) that secured a social target at the center of one side of the box. Each test mouse was placed at the center of the open field and permitted to move freely for 150 seconds in both the absence and presence of a social target (i.e., an unfamiliar CD-1 mouse). All mouse behaviors were recorded via a digital video camera under infrared light illumination. The time spent in the interaction zone (8 cm from the mesh cage) was analyzed using EthoVision XT (Noldus, Netherlands). A social interaction ratio was computed as the ratio of time spent in the interaction zone in the presence and absence of the target CD-1 aggressor mouse. Resilient mice were defined as those with interaction ratios >= 1, whereas susceptible mice were those with interaction ratios < 1.

For a more detailed comparative analysis, mice were further categorized into extreme and mild groups. The extreme group consisted of the 10 most resilient mice with the highest interaction ratios and 10 most susceptible mice with the lowest interaction ratios, whereas the mild group consisted of the 8 resilient mice with the lowest interaction ratios and 8 sensitive mice with the highest interaction ratios.

### 3D pose estimation

To detect the 3D coordinates of each body part of mice, we used the recently developed CNN algorithm, AVATARnet (Kim et al., 2022), which converts multi-view images from the five cameras during the open field test into 2D coordinates of nine body points – nose, head, torso, right and left front limb, right and left hind limb, anus, and tail – in each image. Each 2D coordinate extracted from the five images was converted into a 3D coordinate system using a previously described algorithm (Kim et al., 2022). Errors and outliers in the resulting 3D coordinate system were manually excluded by reconstructing the 3D coordinates into an action skeleton.

### Conventional behavior analysis

The general movement of mice during the OFT was examined by focusing on the center point between the mouse’s head and torso utilizing the 3D coordinates extracted from AVATARnet. Anxiety-like behavior in mice was assessed by characterizing the percentage of time spent and distance moved in a virtual center zone (6 × 6 cm) delineated in the middle of the AVATAR studio chamber. Distance traveled was calculated as the cumulative length of the center point’s movement throughout the experimental period. Rearing behavior was defined as movement that caused the height of the center point to exceed 1.5 times the mean height. These conventional mice behavior parameters were validated by experimenters.

### Machine learning classification of motion sequences and mice

The linear SVM decoder was trained on the continuous 3D coordinates by first dividing the 400 seconds of 3D-coordinate data into specific time durations. The resulting subdivided 3D coordinates were flattened into what is termed a motion sequence. Each motion sequence was labeled as resilient or susceptible and divided into training and test data sets at a ratio of 80:20. The number of mice in resilient and susceptible groups was equalized (n = 15); for susceptible mice (n = 28), this entailed randomly selecting 15 mice for each training/test iteration. The performance of the SVM decoder was then compared by training it on different lengths of motion sequences, from 1 to 10 seconds in length. This analysis identified a 4-second motion sequence as showing the highest decoding performance in classifying the motion sequence. The performance of the SVM decoder was evaluated by comparing the outcomes of 100 bootstrap iterations of random hold-out cross-validation with SVM trained with a random-shuffled dataset by t-test. All motion sequence scores evaluated by the SVM (100 motion sequences during the OFT) were averaged, and mice with an average score > 0.5 were classified as resilient.

### Identification of behavior features

Characteristic features were identified by performing supervised dimensionality reduction and unsupervised clustering (William et al., 2022). Behavioral data from highly resilient and highly susceptible mice, which showed better decoding performance, were used for these analyses. Motion sequences in these groups that were correctly classified by the SVM decoder were used to capture each group’s most significant behavior features. Using a rotation matrix (rotation transform), we then pre-processed motion sequences so as to have the same starting quadrants in the test chamber while conserving their position in each quadrant. This pre-processing of 3D pose data was found to be essential as it removed the bias effect of the starting position of each motion sequence in the subsequent dimensional reduction process.

A supervised dimensionality reduction approach (PLS), was then applied to further reduce the dimensionality of the data using a built-in function in MATLAB. K-means clustering, employing the optimized cluster number found by the MATLAB built-in function, evacluster, which uses the DaviesBouldin clustering algorithm, was used for behavioral clustering. We confirmed that the captured behavioral features could be generated in mice not used in behavior identification, reducing the possibility of capturing overfitted characteristics in the data (Fig. 4). The representative behavior in each motion sequence cluster from k-means clustering was identified by selecting the centroid of each cluster. Thereafter, the 30% of motion sequence points closest to the centroid in each cluster in the PLS dimension were selected and the motion sequence was reconstructed into a 3D coordinate system and action skeleton using a previously described, open-source algorithm (https://actnova.ngrok.io/).

Each of the cluster-like behaviors was quantified in each mouse by training the one-class SVM decoder on the specific cluster’s representative behavior and the other clusters’ representative behaviors using a built-in function in MATLAB. The number of both representative behaviors of the specific cluster and those of the other clusters was equalized by randomly selecting other clusters’ representative behaviors for repeated iterations covering all datasets, as was done for k-fold cross-validation. After training, all motion sequences of each mouse during the OFT were evaluated by the SVM decoder, and the total score, evaluated as the specific cluster-like behavior in each mouse, was summed. The performance of one-class SVM decoding was confirmed using k = 8-fold cross-validation (average per cluster, 84%). The cluster-like behavior numbers characteristic of resilient and susceptible mice were compared using a t-test, and their correlations with actual SI ratios were determined using Pearson’s correlation coefficient.

### Prediction of SI ratio using GLM

The logistic GLM was fitted from the quantified number of resilient and susceptible cluster-like behaviors using a built-in function in MATLAB. The score predicted by the GLM was then compared between resilient and susceptible mice using a t-test, and its correlation with the actual SI ratio was calculated using Pearson’s correlation coefficient.

## Notes

### Competing Interest Statement

The authors have declared no competing interest.

