## Extended Data for "Back-to-the-future motion analysis using machine intelligence predicts the potential risk of mood disorders"

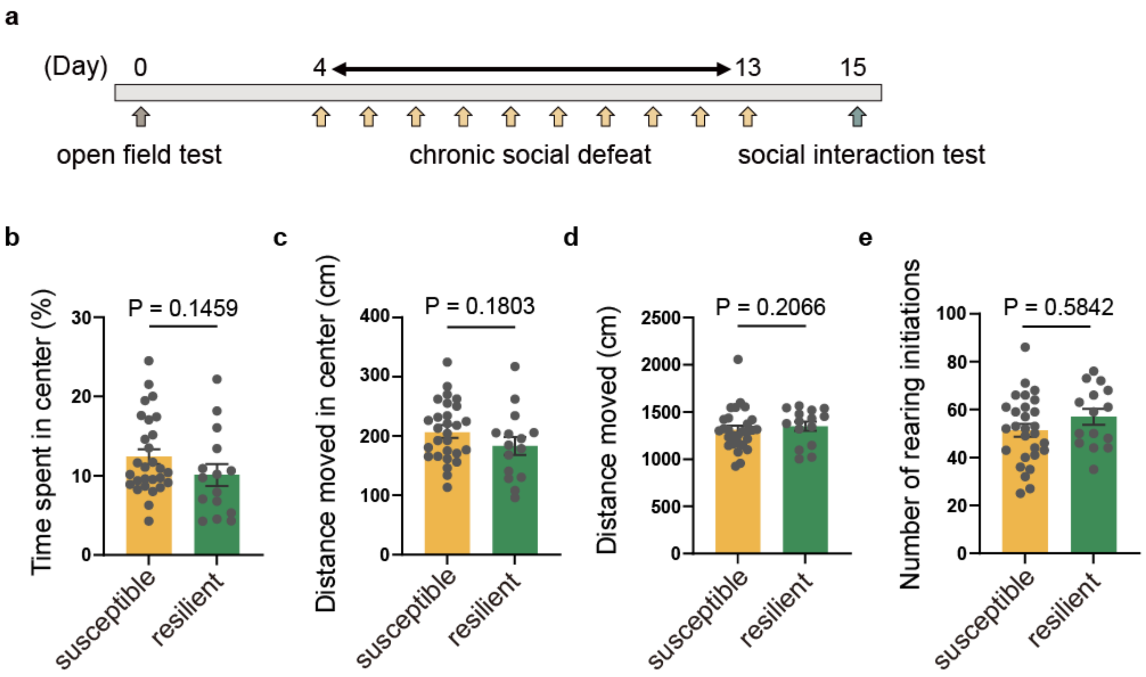


Extended Data Fig.1: Experimental timeline and conventional behavioral analysis in an OFT before CSDS.

a, Experimental timeline. b, Comparison of the relative time spent in the center zone in an OFT between susceptible and resilient mice prior to the social defeat stress protocol. Data are presented as means ± s.e.m., where each dot represents an individual mouse (susceptible, n = 28; resilient, n = 15; Mann-Whitney test). c–e, Comparison of the distance moved in the center zone (c), total distance moved (d), and rearing initiation (e) between susceptible and resilient mice (unpaired t-test).

Cluster 1 (R 58.02%) : Rearing followed by crouching

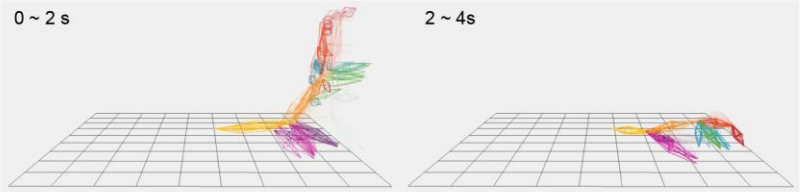


Cluster 2 (R 57.75%) : High and long rearing

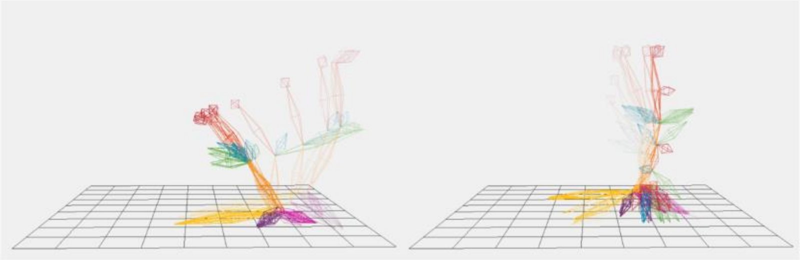

Cluster 3 (S 51.47%) : Dart (moving with high acceleration and stopping)

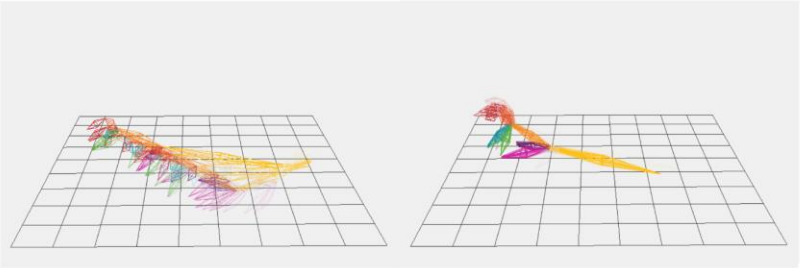

Cluster 4 (R 58.67%) : Rearing and moving toward the corner

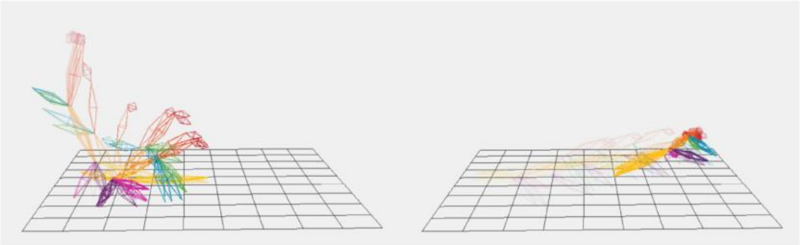

Cluster 7 (S 55.92%) : Circling (rotating in a fixed position in bending body)

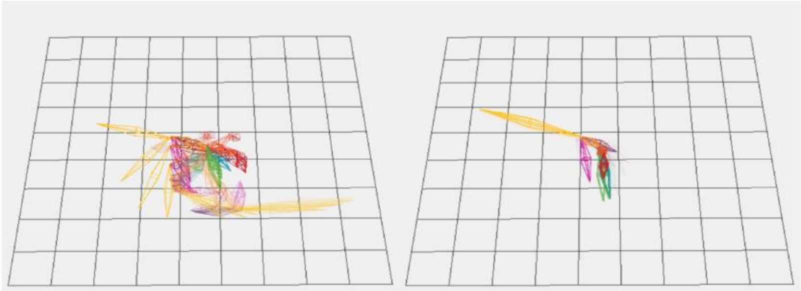

Clsuter 9 (S 51.68%) : Foraging and circling

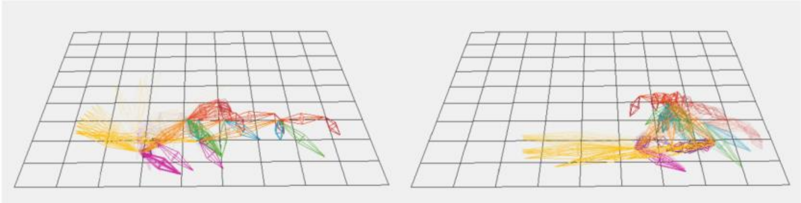

Extended Data Fig.2: Representative behavior of each motion sequence cluster.

Reconstructed 3D motion sequences of each cluster. The 4-second motion sequences are divided into two capture figures (0–2 and 2–4 seconds). The transparency of motion sequences denotes sequential changes over time.

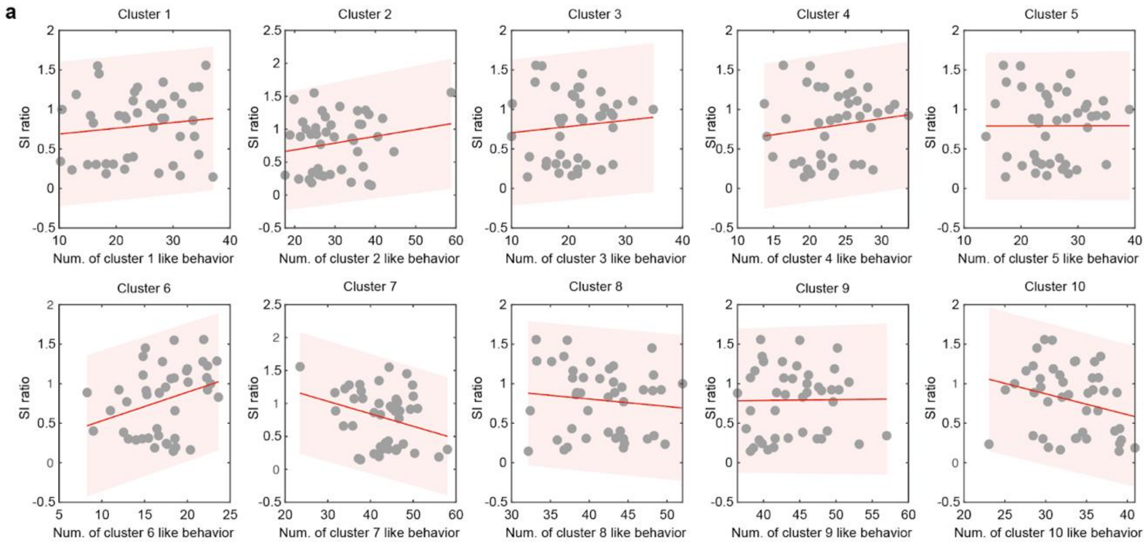


Extended Data Fig.3: Correlations between SI ratio and the frequency of behaviors in each identified behavior cluster in all mice.

a, Quantification of the linear correlation between the SI ratio and the frequency of behaviors in each identified behavior cluster. Pearson's correlations: cluster 1, 0.1255; cluster 2, 0.1944; cluster 3, 0.1002; cluster 4, 0.1487; cluster 5, 0.0017; cluster 6, 0.3186; cluster 7, -0.2987; cluster 8, -0.1118; cluster 9, 0.0115; cluster 10, -0.2626.

Supplementary Video

https://www.youtube.com/watch?v=k-LRbzhag8c
